# Environmental and evolutionary drivers of the modular gene regulatory network underlying phenotypic plasticity for stress resistance in the nematode *Caenorhabditis remanei*

**DOI:** 10.1101/243758

**Authors:** Kristin L. Sikkink, Rose M. Reynolds, Catherine M. Ituarte, William A. Cresko, Patrick C. Phillips

## Abstract

In response to changing environmental conditions, organisms can acclimate through phenotypic plasticity or adapt by evolving mechanisms to cope with novel stressors. Changes in gene expression, whether dynamic or evolved, are an important way in which environmental responses are mediated; however, much is still unknown about how the molecular networks underlying plastic phenotypes evolve. Here, we compare transcriptional responses to acute heat stress among four populations of the nematode *Caenorhabditis remanei*—one selected to withstand heat stress, one selected under oxidative stress, an unselected control, and the ancestral population. We used a weighted gene coexpression network analysis within these lines to identify transcriptional modules, which are sets of genes that respond similarly to stress via plastic responses, evolutionary responses, or both. The transcriptional response to acute heat stress is dominated by a plastic response that is shared in the ancestor and all evolved populations. However, we also identified several modules that respond to artificial selection by (1) changing the baseline level of expression, (2) altering the magnitude of the plastic response, or (3) a combination of the two. Our findings reveal that while it is possible to perturb the nature of the transcriptional response network with short bouts of intense selection, the overall structure of transcriptional plasticity is dominated by inherent, ancestral regulatory systems.

## INTRODUCTION

When faced with novel and stressful environmental conditions, individual organisms must be able to acclimate in order to survive, and populations of organisms will often need to adapt to flourish in the new conditions (Hoffmann and Hercus 2000). The coherent induction of novel trait values via phenotypic plasticity in response to environmental variation is one mechanism by which organisms can increase their fitness when faced with an environmental challenge (Bradshaw 1965). Like other complex phenotypes, phenotypic plasticity has a genetic basis, and therefore can evolve in response to selection (West-Eberhard 2003; Moczek *et al*. 2011). The adaptive response of a population to new, stressful conditions may hence involve the evolution of novel patterns of phenotypic plasticity (Via and Lande 1985; Gomulkiewicz and Kirkpatrick 1992; Gavrilets and Scheiner 1993; Lande 2009; 2014). Furthermore, adaptation to novel environments in the wild may require the change of myriad characters in response to numerous stresses, leading to a potential correlated response in mean phenotypes in one environment (e.g., Grant and Grant 1995; Fischer *et al*. 2007). Covariance in patterns of phenotypic plasticity of different traits across environments (e.g., Czesak *et al*. 2006; Stinchcombe *et al*. 2010) can also evolve if the plastic responses share a genetic basis.

Plasticity has been studied in the laboratory and the field for more than a century at the phenotypic level (Baldwin 1896b; 1896a; Clausen *et al*. 1940; Waddington 1953; 1956; Schmitt *et al*. 1995; Bennett and Lenski 1997; DeWitt 1998; Nussey *et al*. 2005; Cheviron *et al*. 2013), and has been shown to be adaptive in many different systems (e.g., Dudley and Schmitt 1996; Agrawal 1998; Aubret *et al*. 2004; Charmantier *et al*. 2008). Despite the long body of work on phenotypic plasticity and its documented importance in adaptation to novel environments, little is known about the molecular basis of plasticity or the adaptive evolution of molecular systems that underlie phenotypic plasticity. While we know that novel environments can induce a phenotypic change that may be adaptive, we do not understand the relative balance between relatively rapid plastic responses and longer term changes in baseline expression of a trait in response to adaptation to a novel environment. Such questions are impossible to answer in most cases because of a confounding of environmental and demographic history in any given natural population. Experimental evolution is therefore an ideal approach to address this problem because proximal exposure to both novel and ancestral environments can be separated from long-term adaptation to the novel environment in a robust and reproducible fashion (Sikkink *et al*. 2015).

While phenotypic plasticity has been studied for a wide variety of traits for decades, the advent of whole-genome functional approaches (e.g. microarrays and RNA-seq) now allow large sets of genes that are differentially expressed in response to particular environmental stresses to be identified (e.g., Gasch *et al*. 2000; Swindell *et al*. 2007; Badisco *et al*. 2011; Schunter *et al*. 2014). Like any trait, plasticity in gene expression can be characterized in terms of its norm of reaction: the change in expression level for a given genotype when it is exposed to one environment versus another (Fig. 1). If populations of organisms continue to experience the new environment long enough for genetic changes to accumulate within the population, then norms of reaction themselves can change, both in terms of basal expression levels and in terms of the plasticity of expression itself (Fig. 1). The advantage of using whole-genome approaches in this context is that the labilities of thousands of genes can be tested simultaneously, allowing both the broad categorization of response profiles and the clustering of genes into distinct response types.

**Figure 1.**
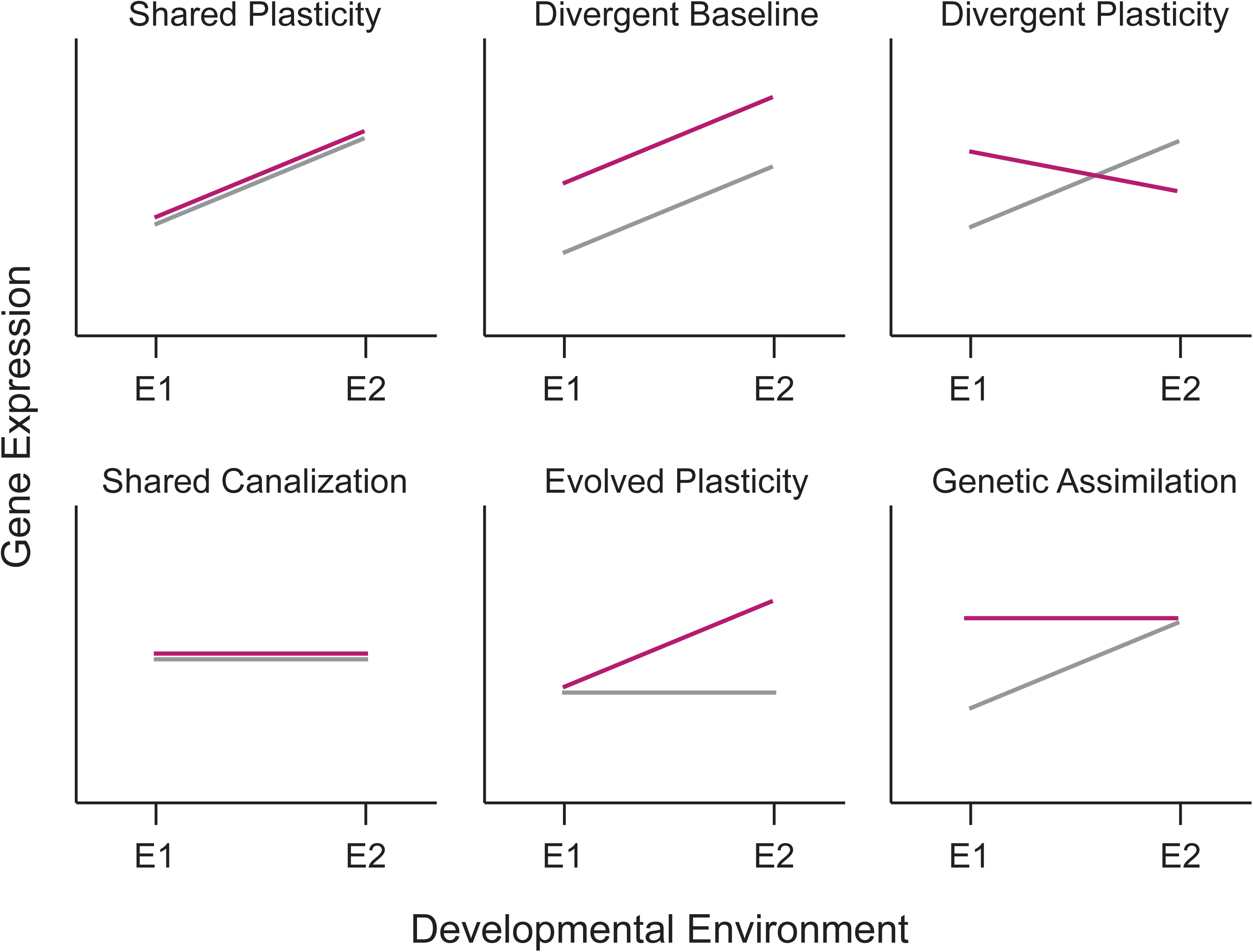
Different patterns for the evolution of phenotypic plasticity in response to environmental change. In each panel, the norm of reaction for the ancestral population is denoted by the gray line and that of the derived population by the red line. Patterns of plasticity can either remain the same, evolve to become different, or change in overall mean level of expression. The various options illustrated here are not an exhaustive list, as combinations of many of these options are also possible.

A central goal of the analysis of gene regulatory networks (GRN) is to identify coregulated sets of genes, which are likely to have similar functions (Eisen *et al*. 1998; Wolfe *et al*. 2005). While network approaches have been applied to understand the basis of plasticity in response to environmental variation (Promislow 2005; Barchuk *et al*. 2007; Rose *et al*. 2016), it is yet unclear how evolutionary changes in GRNs contribute to the evolution of phenotypic plasticity itself. Performing systems analyses of plasticity within an experimental evolutionary context provides a powerful opportunity to simultaneously identify the loci or pathways that are responsive to environmental perturbation, as well as the nature of evolution within those pathways over time. Here we use a GRN analysis approach to understand the evolution of phenotypic plasticity in experimentally evolved populations of nematode worms.

We evolved populations of the nematode *C. remanei* in the laboratory, selecting for resistance to heat stress and oxidative stress under several environmental conditions (Sikkink, Reynolds, *et al*. 2014; Sikkink et al. 2015). As a result of selection, most populations exhibited changes in phenotypic plasticity across environments (Sikkink, Reynolds, *et al*. 2014; Sikkink *et al*. 2015). Here we used a differential gene expression approach via RNA-seq to determine the structure and evolution of the gene coexpression network in a set of populations derived under this experimental framework. We used deep and highly replicated sampling to obtain transcriptional profiles of ancestral and stress-selected populations under multiple environmental conditions. We reconstructed the coexpression network to identify transcriptional modules associated with the plastic response to stress and/or the evolution of that response. We also used this analysis to assess whether adaptation to stress involved the same co-regulated modules—and therefore likely the same pathways—in lines selected to withstand different stressors. The contrast between environmentally induced changes in gene expression between ancestral and evolved populations allows the identification of the primary modes of expression-based responses to stress, as well as a contrast between extant and evolved plasticity in determining adaptation to a novel environment.

## MATERIALS AND METHODS

### Experimental evolution of *C. remanei*

We used the experimentally evolved populations of *C. remanei* that have previously been described by Sikkink *et al*. (Sikkink, Reynolds, *et al*. 2014; Sikkink, Ituarte, *et al*. 2014; Sikkink *et al*. 2015). Briefly, 26 isofemale strains of *C. remanei* were isolated from terrestrial isopods (Family *Oniscidea)* collected from Koffler Scientific Reserve at Jokers Hill, King City, Toronto, Ontario. These strains were crossed in a controlled fashion to promote equal genetic contributions from all strains. The resulting genetically heterogeneous population (PX443) was the ancestral population for the experimental evolution.

A subset of the ancestral population was used for transcriptional profiling. In addition to the ancestor, three experimentally evolved populations were sampled for RNA-sequencing. All selection lines had been evolved at 20°C as described previously (Sikkink, Reynolds, *et al*. 2014; Sikkink *et al*. 2015). One representative control population, one heat-selected population, and one oxidative-selected population were used. The heat-selected line was generated by exposing age-synchronized L1 larval worms to a 36.8°C heat shock approximately every second generation. The oxidative-selected was similarly treated with a 1mM solution of hydrogen peroxide. The control populations received a mock selection treatment, from which worms were selected at random to continue the selected line at a similar census size. All lines were frozen after every two selection events. The final experimentally evolved populations used for the transcriptomics had experienced a total of 10 acute selection events and five freeze-thaw cycles.

### Expression data

We collected L1 tissue from the ancestral, control, heat-selected, and oxidative-selected populations to use for transcriptional profiling (Supplemental Fig. S1). All lines except the oxidative-selected population have been previously described (Sikkink, Reynolds, *et al*. 2014; Sikkink, Ituarte, *et al*. 2014). Briefly, we thawed frozen stocks of worms from each population. Except in the oxidative-selected population, 6 replicates per treatment were collected from a minimum of two independently thawed populations from each line. For the oxidative-selected line, all replicates were collected from a single thawed population of worms. Worms were raised at 20°C until the population was large enough to collect enough individuals for RNA isolation. Age-synchronized L1 larvae were raised for 20 hours in liquid medium at either 20°C or 30°C (Supplemental Fig. S1). Prior to tissue collection, larval worms were passed through a 20-μm Nitex screen to remove unhatched eggs and dead adults. Total RNA was isolated from approximately 100,000 pooled individuals using standard TRIzol methods. Sequencing libraries were prepared according to the protocols as previously described (Sikkink, Reynolds, *et al*. 2014; Sikkink, Ituarte, *et al*. 2014). Samples were sequenced from a single end, to a length of 100 nucleotides in six lanes on an Illumina HiSeq 2000 at the University of Oregon Genomics Core Facility.

### Initial processing of RNA-seq data

Initial quality filtering of raw sequence reads was performed using the *process_shortreads* component of the software Stacks (Catchen *et al*. 2011; 2013). Reads were discarded if they failed Illumina purity filters, contained uncalled bases, or if sample identity could not be determined due to sequencing errors in the barcode sequence. Reads with ambiguous barcodes were recovered if they had fewer than two mismatches from a known barcode. Using the alignment software GSNAP (Wu and Watanabe 2005; Wu and Nacu 2010), we aligned all reads that passed the quality filters to a reference genome for *C. remanei* assembled from strain PX356 (Fierst *et al*. 2015). We then used the htseq-count tool from the Python package HTSeq (Anders *et al*. 2015) to count all reads unambiguously aligning to gene models.

For all expression analyses, we first normalized the gene counts from all samples to account for differences in library size using the scaling procedure implemented in the *DESeq2* package (Anders and Huber 2010; Love *et al*. 2014) in R (R Development Core Team). The expression dataset was next filtered to exclude the 40% of genes with the lowest variance across treatments. Independent filtering of genes with very low variance in expression across treatments generally improves power in subsequent analyses (Bourgon *et al*. 2010; Anders *et al*. 2013). The expression values of the remaining genes were transformed using the variance-stabilizing transformation within *DESeq2* for further analysis.

Finally, we used surrogate variable analysis (SVA) to reduce signal from batch effects and other unknown sources of variance (Leek and Storey 2007). SVA uses the residual expression matrix after accounting for the variables of interest—in this case temperature and population—to identify latent variables within the expression matrix. Using the R package *sva* (Leek and Storey 2007), we identified five latent variables encapsulating the unknown effects. To remove the effects of these variables from the gene expression data, we used *limma* (Ritchie *et al*. 2015) to build a regression model including only the latent variables obtained from SVA. We retained the residual expression matrix for all downstream analyses.

### Multivariate analysis of transcriptional variation

We used non-metric multidimensional scaling (nMDS), which is an unsupervised ordination method that enables highly-dimensional data to be projected onto a few axes for visualization. For RNA-seq data, nMDS may preferable as an ordination method, because it does not assume linear relationships within the data, enabling nMDS algorithms to robustly extract complex patterns from gene expression data (Taguchi and Oono 2005). One drawback of this nonparametric approach is, however, that the scores for variables mapped onto ordination axes can not be easily interpreted (in contrast to principal component scores, for example), and other methods may be required to identify genes contributing to differences between groups.

To carry out the nMDS ordination, a dissimilarity matrix was calculated for the filtered dataset of SVA residuals using Bray-Curtis dissimilarities (Bray and Curtis 1957). Using other distance metrics did not substantially alter the ordination plot. Data transformation, ordination, and scaling were performed in five dimensions using the *vegan* package (Oksanen *et al*. 2013). We tested for significant differences among populations and treatments using a permutational analysis of variance performed on the Bray-Curtis dissimilarity matrix. Population, treatment, and the interaction term were included as effects in the model, and 1000 permutations were run.

### *de novo* Network Analysis

We used weighted gene coexpression network analysis (WGCNA), implemented in the R package *WGCNA* (Langfelder and Horvath 2008) to identify sets of genes (modules) that are highly correlated in their expression patterns. The initial network was constructed from all replicate samples for all treatments (n=48) using a signed adjacency matrix with power = 5 to construct the topological overlap matrix. Hierarchical clustering of the topological overlap matrix was performed using the hclust function of flashClust (method = “average”) (Langfelder and Horvath 2012). Initial module assignments were made using the dynamicTreeCut algorithm (Langfelder *et al*. 2008) with the following options: cutHeight = 0.905, deepSplit = 2, minClustSize = 30, pamRespectsHybrid = FALSE. We used a resampling approach to determine the probability that each gene was assigned to the appropriate module. To do this, we selected four of the six replicates for each treatment group at random to create a new subsampled dataset with 32 samples each. A total of 100 resampled datasets were created in this way. Using the same parameters as in the full dataset, we reconstructed the gene coexpression network for each of the resampled datasets. We then assessed whether each gene belonged in a given module by the following criteria: (1) If a module within a resampled network comprised at least 10% of a module in the full network, the genes in the resampled module were considered a significant group within the original module. Thus, each module from the full network could consist of several well-supported modules from the resampled data, but the network topology is biased toward the modules created from the full dataset. (2) Every gene from the resampled modules that was included in such a group in at least 70% of the resampled networks was determined to be strongly supported as a member of that module in the original network. All genes that did not meet these criteria were removed to the “unassigned” bin. After poorly supported genes were removed form each module, we merged highly correlated modules together based on the correlations among module eigengenes. Modules with highly correlated eigengenes (r > 0.9) were merged into single modules. Finally, the genes from any remaining modules that contained fewer than the minimum cluster size of 30 genes were moved to the “unassigned” bin.

### Functional Annotation of Evolving Modules

The eigengene for each module, defined as the first principal component of the expression of all the genes in the module, was calculated to represent the general pattern of expression seen within each module. With the samples from the ancestral population, we first performed a t-test for significant differences in eigengene expression across the two environments to identify modules contributing to the ancestral response to thermal stress. We then performed an analysis of variance on module eigengenes to test for effects of population, temperature, and population-by-temperature interactions on the overall module expression. For modules that had a significant effect of population, we used Tukey HSD to identify pairwise differences between lines. All statistical tests on the eigengene expression were carried out in the R statistical environment (R Development Core Team).

To functionally annotate the genes expressed in our data, we searched the nr protein database using BLAST+ (v2.2.28) (Camacho *et al*. 2009) to find significant (E-value > 10^−3^) matches to each gene. We used Blast2GO v3.0 (Conesa *et al*. 2005; Conesa and Götz 2008) to map significant BLASTx results to gene ontology terms and to compute a Fisher’s Exact Test to test for significant over-representation of GO terms in each module.

We examined each module for enrichment of known regulatory targets of 23 transcription factors for which binding data is available for the related nematode *C. elegans*. Binding targets for all transcription factors except for the FOXO transcription factor DAF-16 were obtained from the *C. elegans* modENCODE project (Niu *et al*. 2011). These targets were all identified from chromatin immunoprecipitation sequencing (ChIP-seq). Putative target genes bound by DAF-16 have been previously identified using two different approaches: ChIP (Oh *et al*. 2006) and DNA adenine methyltransferase identification (DamID; Schuster *et al*. 2010). Target genes could be included multiple gene sets if they are bound by more than one transcription factor.

*C. remanei* homologs for each of the *C. elegans* transcription factor targets were determined based on the annotations that have been curated in the WS220 release of WormBase (Harris *et al*. 2009). Homologous genes identified by any method were included as possible transcription factor targets in *C. remanei*. In cases where multiple *C. remanei* genes were matched to a single gene in *C. elegans,* all possible homologous genes were included in the gene set, since no information was available to determine whether transcription factor binding was preserved preferentially in either possible homolog.

Modules were tested for significant enrichment of target genes bound by each transcription factor using a one-tailed Fisher’s exact test. In addition, we tested for enrichment of the *C. remanei* heat shock proteins previously identified (Sikkink, Reynolds, *et al*. 2014).

### Data Availability

Sequence data for the ancestral, control, and heat populations were previously deposited in the NCBI Gene Expression Omnibus (GEO) database as part of series GSE56510 with accession numbers GSM1362987-1363022. Additional RNA-seq data for the oxidative population were deposited in GEO with the accession number XXXXXX.

## RESULTS

### Divergence occurs in transcriptional regulation across temperatures and between evolved populations

We first sought to determine whether samples from the various populations or temperatures could be differentiated based on global patterns of gene expression. To do this, we used non-metric multidimensional scaling (nMDS), a powerful ordination method that does not assume linear relationships among variables (Taguchi and Oono 2005). On the first axis, we observed distinct separation between the two temperature treatments (Fig. 2A). To assess the significance of the observed differences between temperature treatments, we used a permutational analysis of variance (PERMANOVA) on the dissimilarity matrix. This analysis confirmed that temperature had a highly significant effect on global gene expression (F_1,40_ = 6.41, *P* = 0.001).

**Figure 2.**
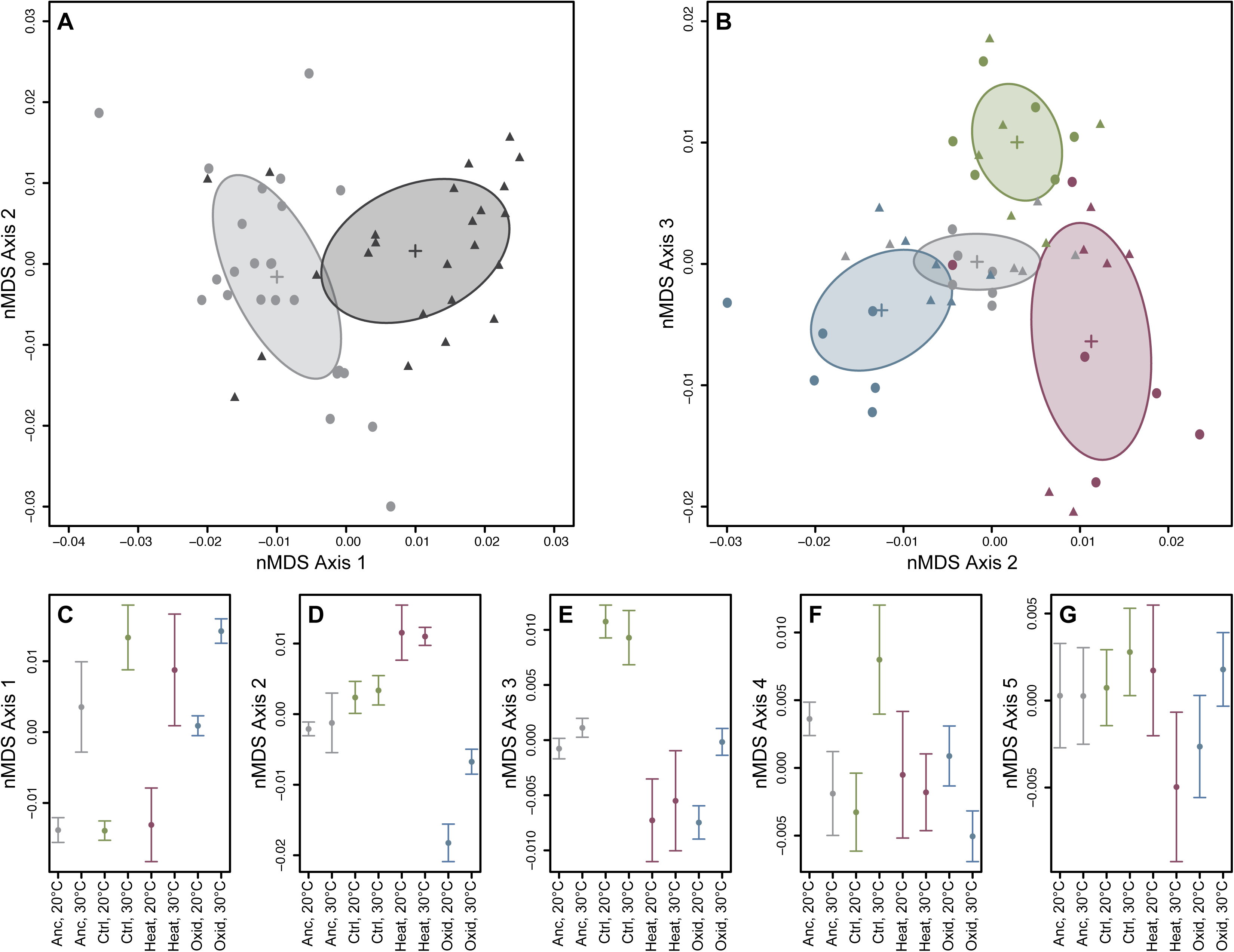
Non-metric multidimensional scaling of RNA-seq samples based on the filtered set of all expressed transcripts. **(A)** Axes 1 and 2 from the ordination. Light grey circles indicate samples raised at 20°C, while dark grey triangles represent samples raised at 30°C. Crosses and ellipses indicate the centroid and standard deviation for each temperature treatment. **(B)** Axes 2 and 3 from the ordination showing the distribution of samples from the ancestor (gray), control (green), heat-selected (red), and oxidative-selected (blue) populations. Circles indicate samples raised at 20°C, and triangles represent samples raised at 30°C. Crosses and ellipses indicate the centroid and standard deviation for each population. **(C-G)** The mean nMDS score (±1 standard deviation) for all treatment combinations on each nMDS axis.

The four populations also differed significantly from one another (PERMANOVA; F_3,40_ = 3.17, *P* = 0.001). The population differences accounted for variation observed on Axis 2 and Axis 3 in the nMDS analysis (Fig. 2B). Both the oxidative and heat selected lines diverged from the ancestral population on Axis 2, but in opposite directions (Fig. 2D). In contrast, the control population separated from the ancestral and selected populations on Axis 3 (Fig. 2E). This pattern of divergence from the ancestor suggests that the three different selection regimes lead to unique changes in transcriptome regulation in these populations. In addition, the response to the temperature treatment was strongly dependent on the population (PERMANOVA, population-by-temperature interaction: F_3,40_ = 1.77, *P* = 0.002). The consequences of the interaction effect are most apparent on Axis 4 (Fig. 2F), where the control-selected population responds to temperature in the opposite direction compared to the remaining lines, and Axis 5 (Fig. 2G), on which the heat population responds in the opposite direction.

### Network modules are differentially associated with line- and temperature-specific variation in expression

Because nMDS is a non-metric method, the contribution of specific genes, or suites of genes, to divergence on each axis is not readily interpretable. To address this limitation, we used weighted gene co-expression network analysis (Zhang and Horvath 2005; Langfelder and Horvath 2008) to identify modules—sets of genes with strongly correlated expression patterns that are more loosely connected to other such modules. We sought to identify modules that were important in the differential regulation of stress resistance in our evolved populations of *C. remanei,* because members of a gene module often share a common function (Eisen *et al*. 1998; Wolfe *et al*. 2005), and highly correlated genes sets may share transcriptional regulators (Allocco *et al*. 2004, but see also Marco *et al*. 2009). Coexpression network analysis can therefore provide unique and useful insights into gene regulatory networks.

Network analysis identified 13 co-expressed modules containing a total of 5,622 genes (Table 1). An additional 9,212 genes were not consistently assigned to any module after resampling and were designated as “Unassigned”. For each module, we calculated the eigengene (Langfelder and Horvath 2008), defined as the first principal component of the module. An eigengene’s expression explains the largest proportion of variance for the genes within the module and is therefore representative of the expression of the combined set of correlated genes within the module (Supplemental Fig. S2). For all assigned (numbered) modules, the eigengene explains more than 40% of the expression variance within the module, with several explaining 50 to 70% (Table 1). The second principal component explains no more than 5.6% in any assigned module (Supplemental Fig. S2), confirming that the eigengene is in fact an appropriate representation of expression for the genes within the assigned modules.

In the balance between environment and evolution in changes in patterns of transcriptional regulation (Fig. 1), plasticity and/or the retention of ancestral plasticity via the evolution of baseline expression predominates. Significant temperature differences or temperature-by-population interactions were observed in 12 out of 13 modules (Table 1). In fact, the two largest modules, Module 1 and Module 2, show effects only attributable to temperature (“shared plasticity”). Module 1 is composed of genes that are upregulated in all populations under immediate heat stress, whereas Module 2 contains genes that are downregulated in response to heat (Fig. 3). The largest class of modules retained the overall pattern of plasticity displayed across populations, but changed their baseline level of expression within each environment (“divergent baseline”, Fig. 3, Table 1). Finally, four of the modules showed heterogeneity among the pattern of plasticity that was dependent upon their specific evolutionary history (“evolved and/or divergent plasticity”, Fig. 3). In one case (Module 5), the oxidative stress population lost its apparent response to heat stress entirely, indicative of genetic assimilation.

**Figure 3.**
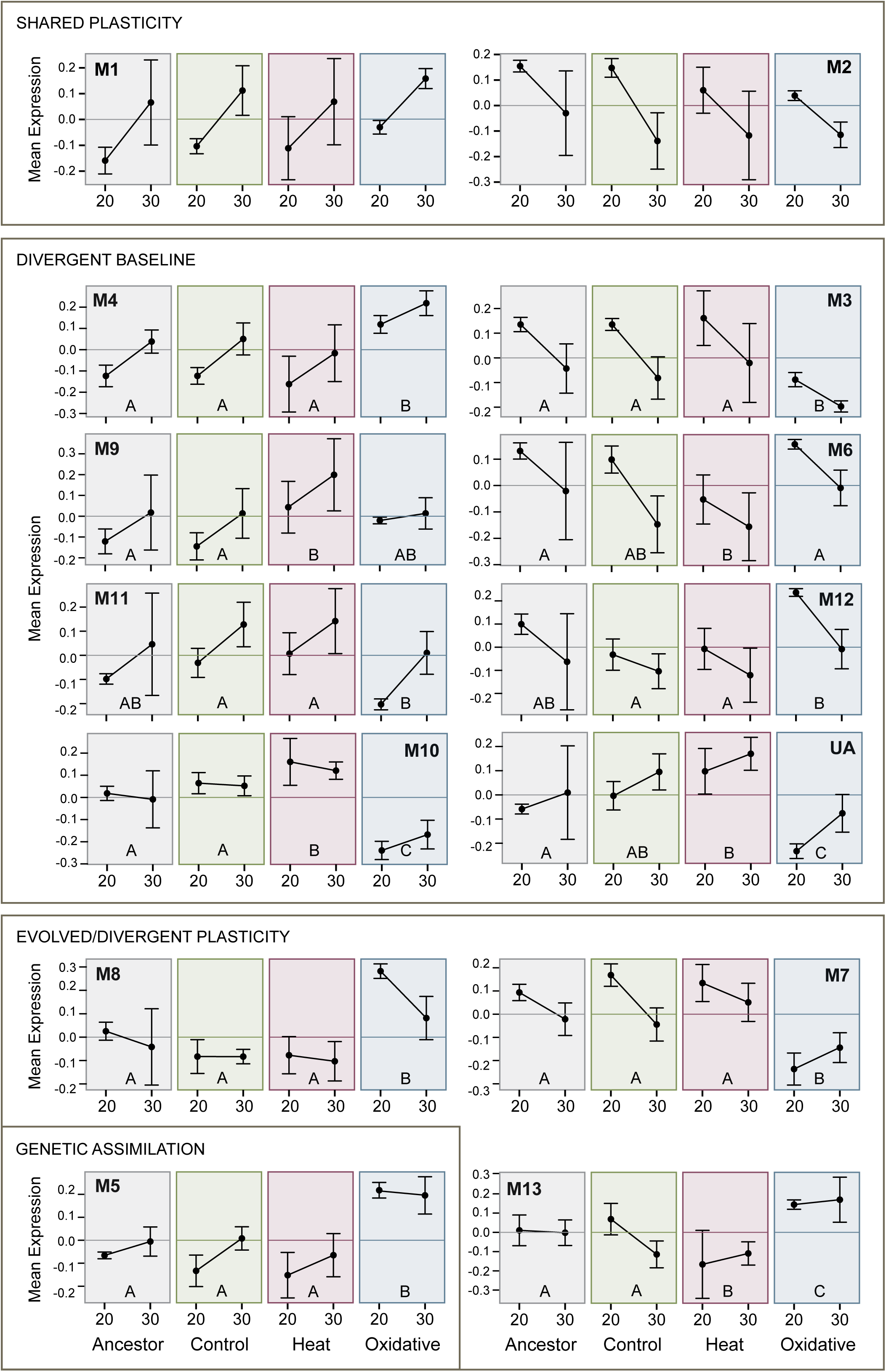
Module eigengene expression across temperatures for each population. Each box illustrates the relative expression for a population (ancestor, control, heat stress, oxidative stress) for individuals raised at either 20° or 30°C. Module ID is specified by the number following “M”. Expression levels for “unassigned genes” are provided under “UA”. Modules are roughly grouped into the categories outlined in Fig. 1. For modules in which there was a significant effect of population (Modules 3-13), the letters indicate populations which were different by Tukey’s HSD. Experimental populations that share a letter are not significantly different in the pairwise comparison.

To more specifically address evolutionary divergence among lines, we examined pairwise differences among them for modules that showed a significant population effect (Fig. 3). Focusing specifically within the ancestral population, five modules (Modules 1, 2, 3, 4, and 7), which together contain 4,836 genes, had significant expression differences across environments (Table 2). Evolved lines that diverged from the ancestral population are of particular interest, as these could indicate a set of genes that are adaptive for stress resistance. Two modules, Module 6 and Module 9, differed in the heat-selected population only. These modules are expected to contain genes that are important for adaptation to heat stress. Similarly, five modules (3, 4, 5, 7, and 8) are significantly different from the ancestor only in the oxidative-selected population. Two additional modules, Module 10 and Module 13, have evolved in both stress-selected populations. In both cases, the responses are in opposite directions in each stress-selected population (Fig. 3), similar to the observed differences on Axis 2 in the nMDS analysis (Fig. 2B).

Surprisingly, among the unassigned genes, we observed a significant effect of both temperature and population in the eigengene expression, supporting the observation that a pattern of “divergent baseline” predominates the overall structure of the evolution of gene expression across the genome (Table 1). Given the conservative approach we used to assign genes into modules following resampling, it is likely that genes contributing to the unassigned modules were falsely removed from a real module. However the proportion of variance explained by the eigengene for the unassigned module is relatively small (9.3%). Therefore, it is unlikely that the lack of module assignment for these genes substantially alters the overall network topology we have observed.

### Gene expression modules are enriched for functionally related genes

We next examined the functional relationships among genes in identified modules by looking for enrichment of Gene Ontology terms within each module, especially terms in the biological process ontology (Fig. 4). Enrichment of molecular function and cellular component terms are shown in Supplemental Figs. S3 and S4, respectively. Modules 1, 4, and 5, which contain genes upregulated in response to temperature, were enriched for terms falling under the GO categories pertaining to embryonic development, reproduction, and cellular transport, as well as metabolic processes relating to nucleic acid metabolism and translation. Module 2, which encompasses the large module of genes down-regulated in response to heat, was enriched for genes involved in cell-cell signaling. Modules 3 and 7, which had lower expression in the oxidative-selected population, were enriched for immune system genes and genes involved with translation, respectively. Modules 9 and 11 were enriched for genes involved in metabolic processes acting on molecules other than nucleic acids.

**Figure 4.**
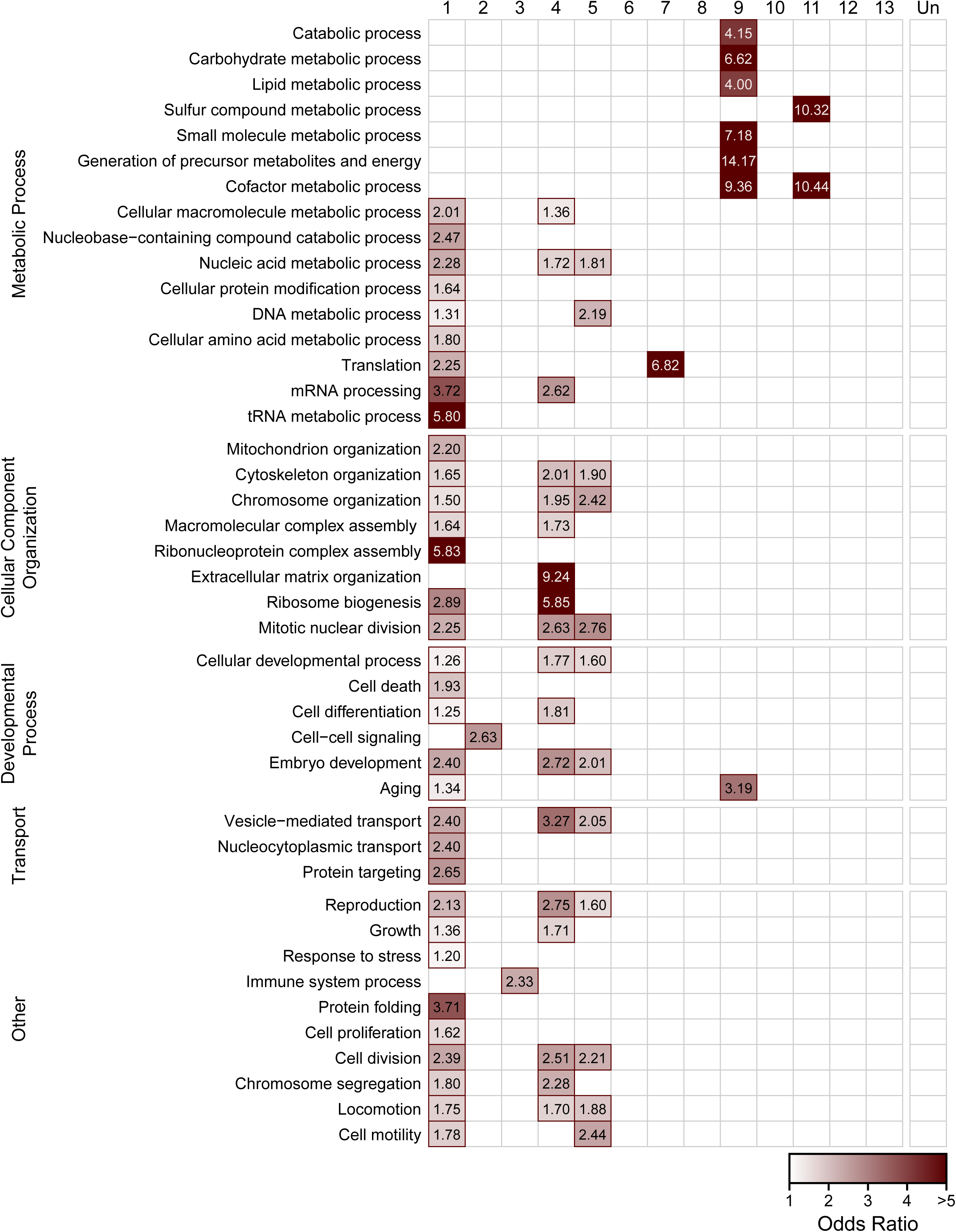
Enrichment of biological process gene ontology terms in coexpression modules. Red outlines and shading signify that there is significant enrichment of genes mapping to the GO term (rows) within a given module (columns) (FDR<0.05). The intensity of the shading corresponds to the odds ratio for the GO term.

### Regulatory targets of stress-responsive transcription factors are enriched in network modules

In *C. elegans,* several transcription factors are known to be critical regulators of cellular responses to stress. However, these regulators may not be differentially expressed in response to stress themselves, but rather undergo protein modifications to activate them under certain conditions. For example, the FOXO transcription factor DAF-16 is a major target of the insulin/insulin-like growth factor signaling (IIS) pathway in worms and is responsible for mediating responses to heat and oxidative stress, among others (Honda and Honda 1999; Hsu *et al*. 2003). DAF-16 is normally localized to the cytoplasm, but in stress conditions, DAF-16 is activated and transported to the nucleus, where it regulates transcription of many target genes (Lin *et al*. 2001; Lee *et al*. 2001). We identified *C. remanei* homologs of known binding targets of 23 transcription factors with previously published transcription factor binding profiles (Oh *et al*. 2006; Schuster *et al*. 2010; Niu *et al*. 2011), and tested for significant enrichment in each of the network modules. We also examined enrichment of another stress-related candidate gene set, the heat shock protein families (*hsps*) previously examined in Sikkink *et al*. (Sikkink, Reynolds, *et al*. 2014).

We observed significant enrichment (FDR < 0.05) of regulatory targets for all but four of the available transcription factors (Fig. 5). Many of the transcription factors are key regulators of embryonic development. Unsurprisingly, Module 1 is enriched for transcriptional targets of most of these genes, including three HOX transcription factors—LIN-39, MBA-5, and EGL-5—consistent with the functional annotation of Module 1’s role in development.

**Figure 5.**
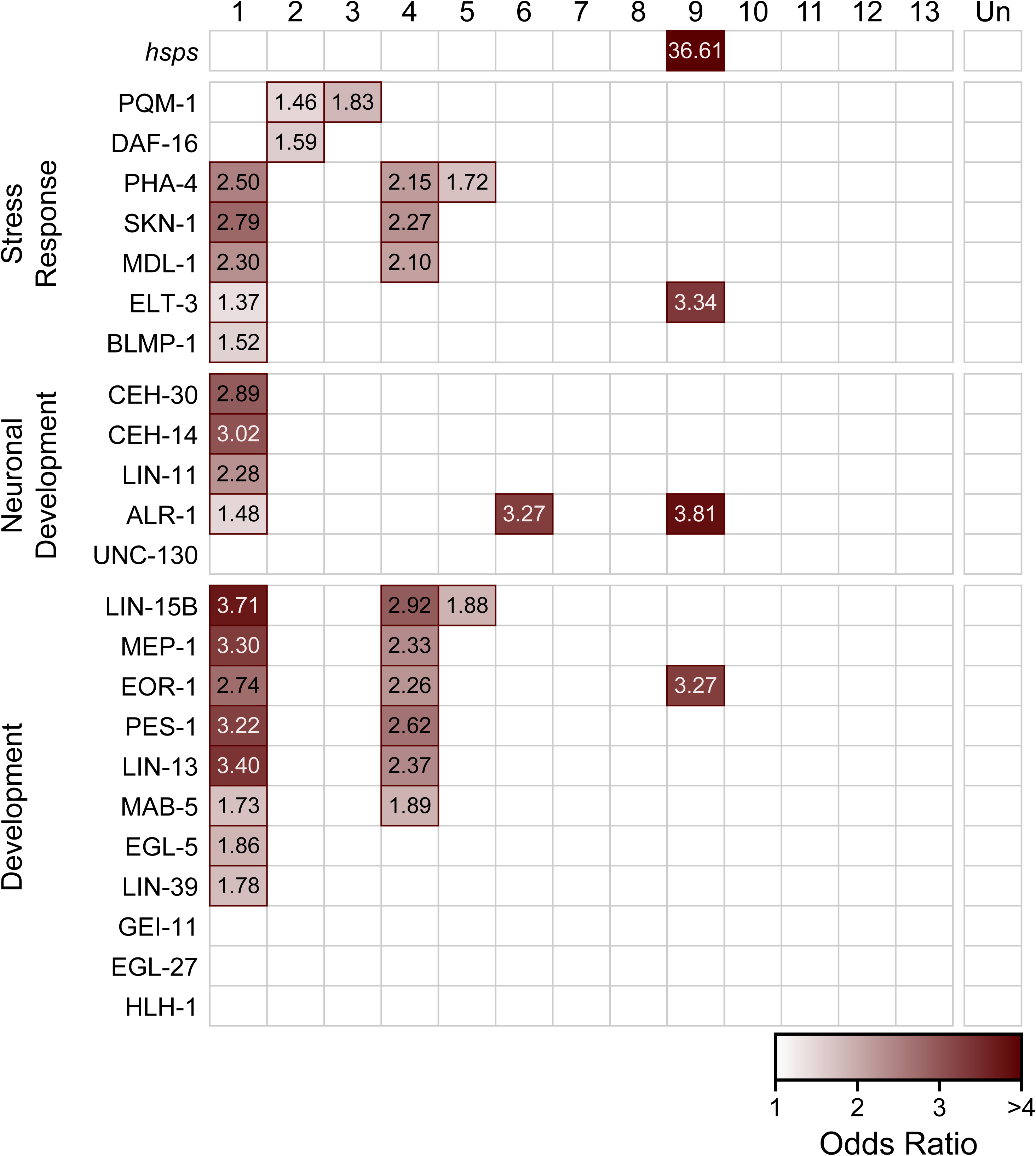
Enrichment of transcription factor target genes in coexpression modules. Each box represents the degree to which the targets of a given transcription factor (rows) are enriched within a given co-expression module (columns). Red outlines signify that there is significant enrichment of target genes in the module (FDR<0.05). Transcription factors are roughly grouped by functional category. The intensity of the shading indicates the odds ratio for the set.

Several transcription factors that regulate stress responses also showed enrichment of their target genes in one or more of these modules. PHA-4, a developmental regulator necessary for formation of the pharynx (Mango *et al*. 1994; Horner *et al*. 1998), has also been implicated in regulating heat shock response through HSP90 (van Oosten-Hawle *et al*. 2013). Targets of PHA-4 were enriched in Module 1 and well as the oxidative-evolved Modules 4 and 5. Genes regulated by SKN-1, another target of IIS that is critical for oxidative stress resistance (An and Blackwell 2003), were also enriched in Modules 1 and 4. Modules 2 and 3 were both enriched for targets of PQM-1, a C2H2 zinc finger and leucine zipper-containing protein (Tawe *et al*. 1998). In *C. elegans,* PQM-1 is responsive to certain types of oxidative stress (Tawe *et al*. 1998), and is a key regulatory target of IIS, in addition to DAF-16 (Tepper *et al*. 2013). Module 2 was also enriched for DAF-16 targets.

Module 9, which was significantly divergent in the heat-selected population, was significantly enriched for heat shock proteins. This module was also enriched for targets of ELT-3, a GATA transcription factor that functions during hypodermal development in *C. elegans* (Gilleard *et al*. 1999) and may also function downstream of IIS to influence longevity (Budovskaya *et al*. 2008), pathogen resistance (Pujol *et al*. 2008), and osmotic stress response (Rohlfing *et al*. 2010). ALR-1, a homeodomain transcription factor involved in development of sensory and GABAergic motor neurons (Tucker *et al*. 2005), and EOR-1 which regulates RAS/RAF-mediated signaling during development (Rocheleau *et al*. 2002; Howard and Sundaram 2002) also bind to more genes than expected within this module.

Modules 7, 8, 10 and 13 were also significantly different between the ancestor and at least one evolved population; however, they did not show significant enrichment of target genes for the available transcription factors. However, ChIP binding data from *C. elegans* was not yet available for some key transcription factors involved in stress response, particularly HSF-1 and HIF-1. Heat shock proteins are known to be regulated by HSF-1 in response to heat stress (Wu 1995; Åkerfelt *et al*. 2010), therefore enrichment of *hsps* in Module 9 may indicate a role for HSF-1 in regulation of that module.

## DISCUSSION

For many organisms phenotypic plasticity is a vital adaptation that has evolved to cope with environmental stress. However, we still know little about the molecular mechanisms contributing to plastic traits. In particular, we know very few of the genes that are involved in phenotypic plasticity, and only a handful of case studies (Promislow 2005; Barchuk *et al*. 2007; Rose *et al*. 2016) have begun to define the genetic regulatory networks that underlie a plastic phenotypic response. Even less is known about how the evolution of phenotypic plasticity is related to the rewiring of gene regulatory networks (GRN) that are environmentally sensitive, particularly when an organism faces adaptation to multiple environments. Here, we examined global changes in a GRN involved in the evolution of phenotypically plastic responses in a powerful experimental evolution system. We used RNA-sequencing and coexpression network analysis to identify sets of coexpressed genes, or modules, which are associated with the evolution of phenotypic plasticity in *C. remanei* in two related, but distinct, evolutionary stresses.

### Global patterns of gene expression describe evolutionary divergence

Based on the global profiles of expression among our filtered set of genes, we observed clear differentiation attributable to the induction of a response to temperature (i.e., plasticity), as well as evolved differences between populations (Fig. 2). Exposure to the inducing temperature resulted in very pronounced changes in the global patterns of gene expression in all populations, which were primarily encapsulated by the primary axis resulting from nMDS analysis (Fig. 2A). The scale of the observed response to the temperature shift in all populations (Fig. 2C) largely confirms our previous observations for the heat selected population—that increased resistance to acute heat stress is not conferred by changing the degree to which genes respond to changes in the thermal environment (Sikkink, Reynolds, *et al*. 2014). However, the more powerful multivariate statistical framework used here reveals that the response to temperature changes did in fact differ among the evolved populations on secondary axes of ordination. These interaction effects were most evident on nMDS Axes 4 and 5 (Fig. 2F and 2G), and suggest that changes in transcriptional plasticity in some parts of the overarching gene regulatory network may be responsible for changes in phenotypic plasticity as well.

In addition to the transcriptional response to temperature, we also detected changes in gene regulation attributable to the evolutionary history of each line. Notably, the three selected populations have diverged from the ancestor in different directions on nMDS Axis 2 and Axis 3 (Fig. 1B). This pattern suggests that at least partially different GRNs contribute to adaptation in each case, likely acting in a modular fashion. These findings are consistent with the observations we have previously made—that there is no genetic correlation between heat and oxidative resistance under the environmental conditions in which these populations evolved (Sikkink *et al*. 2015). In short, although one might reasonably hypothesize a correlated selective response to heat and oxidative stresses that acts through a generic stress response pathway, our data support the alternative hypothesis that evolution results from changes in different GRNs, or least different modules within a GRN, for these two related stresses.

### Modularity of stress GRN evolution

The pattern of expression differences that we observed in our data indicates a high degree of modularity within the gene regulatory network. Despite a relatively small number of experimental treatments, we were able to identify 13 transcriptional modules with highly correlated patterns of expression. Furthermore, the eigengenes that describe expression patterns within each module are differentially associated with the experimental treatments.

A test of our ability to draw meaningful inferences from our RNA-seq data is to examine a well-known pathway. Heat shock proteins *(hsps)* are molecular chaperones known to be a critical component of response to heat stress (Lindquist and Craig 1988). Therefore, we expect these genes to form one or more modules that covary strongly with temperature. Module 9 seems to fulfill this expectation by capturing many of the expected elements of the *hsp* response. This module was strongly enriched for the set of heat shock proteins (Fig. 5), and was also significantly regulated by temperature (Table 1). GO enrichment analysis indicated that this module was strongly enriched for genes involved in a number of metabolic processes, as well as genes involved in aging, although surprisingly “stress response” was not enriched in this module.

Within this heat shock response module, significant expression differences attributable to line were observed (Fig. 3). On closer examination, the heat-selected population was the only selected line to show divergence from the ancestral population. The eigengene expression of this module reveals that the heat-selected population has higher overall expression of these genes relative to the other populations even at 20°C (Fig. 3). However, these genes were still upregulated in response to temperature. The shift in expression in this module is consistent with our previous observation that the heat selected line has evolved resistance to heat stress by shifting the thermal threshold for induction of the heat stress response (Sikkink, Reynolds, *et al*. 2014).

Most of the identified modules, with the exception of Modules 1 and 2, show significant evidence for an evolved response in either of the stress-adapted populations (Table 1) and are of particular interest as candidates to fulfill this role. Interestingly, the response to evolutionary pressure in the heat and oxidative selected lines occurs primarily in independent modules, consistent with the lack of phenotypic correlated responses we have observed previously in this system (Sikkink *et al*. 2015).

Two modules, Module 10 and Module 13, do show evidence of evolutionary change in regulation in both the heat- and oxidative-selected lines (Table 1). However, our data do not support the evolution of a generalized stress response pathway contributing to adaptation in both of the populations that were evolved under these different stressors. In both modules, selection for heat resistance results in an overall change in gene expression in one direction, while selection for oxidative resistance occurs in the opposite direction (Fig. 2). Presumably, these modules would contain a portion of the overlap in pathways expected based on the pleiotropy in the known stress response networks in *C. elegans* (Sikkink *et al*. 2015). However, these two modules together contain only 104 total genes, about 5% of the genes assigned to evolved modules, and the effect of selection to one stressor is antagonistic to the preferred response for the other stress. The overall independence we observed in the genetic network here may contribute to the decoupling of the phenotypic responses to these different stressors, as certain modules within the greater response network have different degrees of pleiotropy, and can potentially allow for fine-tuning of the response.

### Adaptation to different stressors involves non-overlapping subsets of the ancestral stress response network

Six modules, together comprising 4836 genes (about 86% of the genes assigned to modules) had significant responses to temperature in the ancestral population (Table 2). Most of these genes were in Modules 1 and 2, which show no evidence for evolved changes in any of the selective populations (Table 1). This suggests that the majority of the ancestral heat shock response remains unchanged under diverse selective scenarios. Several of the modules that have evolved only in the oxidative selective environment (Modules 3, 4, and 7) also had a significant response to temperature in the ancestral population (Tables 1 and 2). A reasonable interpretation of this pattern is that these modules contributed to the core stress response in the ancestral population, along with the genes in Modules 1 and 2. However, in the oxidative-selected populations, different subsets of genes within the GRN are selected upon, enabling resistance to the oxidative stress instead of heat stress.

In contrast, in the modules that respond to heat selection (Modules 6, 9, 10, and 13), there was no strong evidence for plasticity in the ancestral population (Table 2). The genes in these modules are therefore unlikely to be involved in the ancestral plasticity in response to the inducing temperature, but rather are other sets of genes from a different regulatory cascade that have been modified under particular conditions. It is surprising that it is the heat-selected line that utilized novel stress response components, while oxidative selection in large part modified the existing stress response pathway. Because the eigengene is a combined metric of expression for multiple correlated genes, a trivial interpretation is that some individual genes within this module do in fact respond significantly to temperature in the ancestor, but we have limited power to detect the change when considered as a group. If this is true, then selection for heat stress may use components of the ancestral heat response, although these modules are still largely independent of those invoked for the oxidative stress response. A more interesting explanation that would explain the pattern is that the core heat stress response in the ancestor is already maximized, and further adaptation therefore requires cooption of additional genes that are not part of the core stress pathway. These interpretations may not be mutually exclusive, and determining the relative contributions of the ancestral vs. novel components of the stress response pathway in adaptation to heat stress will require additional investigation into the roles of individual genes within each population.

### Regulation and function of the evolved plasticity GRN modules

Genes that are co-regulated by a common transcription factor are likely to have highly correlated expression (Marco *et al*. 2009) and therefore should be classified as part of the same module. Identifying the transcriptional regulators of each module can provide important insight into which pathways contribute to the evolution of plasticity. In this study, we tested for enrichment of known targets of 23 transcription factors within each of the identified gene modules (Fig. 4). Most of these transcription factors have vital roles in regulating developmental processes, but a few also have well-characterized roles in mediating stress responses to a variety of different stress types.

Many of the tested transcription factors are enriched in Module 1 (Fig. 4). Given that the tested factors are key regulators of development, it is not surprising this module is also functionally annotated as involved in growth, embryonic development, and reproduction— organismal processes which are highly dependent on temperature in the poikilothermic *C. elegans* and its relatives (Riddle 1997). Consistent with that role, the genes in Module 1 do not appear to have evolved differences in gene expression between the ancestor and any of the evolved populations, and likely represent a core set of highly conserved developmental programs that are difficult to alter on short evolutionary timescales.

Not all of the components of these pathways are so highly conserved however. Module 4, for example, is also enriched for regulatory targets of many of the same transcription factors as Module 1, with the notable exception of the transcription factors required for neuronal developmental (Fig. 4). This module had higher overall expression in the oxidative-selected population compared to the ancestor. It is reasonable to think that tissue-specific differences and the combinatorial control of gene expression might allow for the increased evolvability of transcription in this module, despite the overlapping functional role of these genes and those in Module 1.

Another intriguing result from this study is the enrichment of PQM-1 targets in Modules 2 and 3. PQM-1, like DAF-16, is a major target of the IIS pathway, and the two transcription factors appear to function in opposition to one another (Tepper *et al*. 2013). Both transcription factors appear to act together to control a portion of the ancestral stress response identified in Module 2. However, Module 3 also shows evidence for evolved differences in expression in the oxidative-selected line (Table 1), and is enriched for targets of PQM-1 (Fig. 4). PQM-1 is known to respond to oxidative stress, although previous studies describe a response to the oxidative stressor paraquat (methyl viologen; (Tawe *et al*. 1998)) rather than the hydrogen peroxide used in this study. Further study will be required to determine whether PQM-1 and its targets are in fact important contributors to evolution of the oxidative stress response.

### Conclusion

We have identified transcriptional modules with patterns of expression consistent with evolutionary response to selection in two different, but related, stress phenotypes. In general, patterns of ancestral plasticity dominated the evolutionary response, with the predominant mode of adaptation being driven by changes in baseline gene expression within an environment rather than changes in plasticity per se. The evolutionary responses to the two different stressors each modified unique modules within the larger transcriptional network, consistent with previous observations of low pleiotropy between these two response types. Surprisingly, the response in the population selected under heat stress conditions did not modify components of the ancestral response to heat shock, while the oxidative selection line did. We identified a number of modules with significant evolutionary and plastic responses that were enriched for targets of key developmental and stress response transcription factors. The functions of these transcription factors are well described in *C. elegans* and *Drosophila,* and provide useful insight for how they might contribute to adaptation in this system, although precise examination of these effects will require additional study. The scale of the total number of genes involved makes this a somewhat daunting task, however. Indeed, a major conclusion from this work is that short-term adaptation, although it may be built upon existing systems that facilitate a plastic response to the environment, can nonetheless alter the overall pattern of regulation of the majority of the genes in the genome (Boyle et al. 2017). Complex responses to environmental stress remain complex even when adapting to simplified selective environments.

## ACKNOWLEDGMENTS

We thank the many volunteers who have assisted with maintaining the *C. remanei* selection lines. This work was supported by a GRFP and DDIG (1210922) from the National Science Foundation to KLS, a Ruth L. Kirschstein NRSA Postdoctoral Fellowship to RMR (AG032900), National Institutes of Health grants to PCP (AG022500, AG043988, GM096008) and WAC (RR032670), and the Ellison Medical Foundation fellowship to PCP.

---

**Table 1.** Eigengene analysis of modules identified in gene coexpression network analysis.

| Module | Number of Genes | % Variance Explained by PC1 | Eigengene Effects | F | df(num),<br>df(denom) | P value | Evolved<sup>a</sup> | Evolved Population | Functional annotation |
| --- | --- | --- | --- | --- | --- | --- | --- | --- | --- |
| 1 | 1992 | 42.4% | Temperature | 46.33 | 1, 40 | **>0.001\*** |  |  | embryo development, reproduction, transport, others |
|  |  |  | Population | 2.55 | 3, 40 | 0.069 |  |  |  |
|  |  |  | Temp x Pop | 0.13 | 3, 40 | 0.942 |  |  |  |
| 2 | 1471 | 42.5% | Temperature | 46.69 | 1, 40 | **>0.001\*** |  |  | cell-cell signalling |
|  |  |  | Population | 2.38 | 3, 40 | 0.084 |  |  |  |
|  |  |  | Temp x Pop | 1.00 | 3, 40 | 0.401 |  |  |  |
| 3 | 865 | 42.3% | Temperature | 48.73 | 1, 40 | **>0.001\*** | ● | ox | immune system process |
|  |  |  | Population | 15.53 | 3, 40 | **>0.001\*** |  |  |  |
|  |  |  | Temp x Pop | 0.85 | 3, 40 | 0.472 |  |  |  |
| 4 | 389 | 49.8% | Temperature | 38.24 | 1, 40 | **>0.001\*** | ● | ox | embryo development, reproduction, transport, others |
|  |  |  | Population | 23.97 | 3, 40 | **>0.001\*** |  |  |  |
|  |  |  | Temp x Pop | 0.48 | 3, 40 | 0.700 |  |  |  |
| 5 | 278 | 50.3% | Temperature | 11.20 | 1, 40 | **0.002\*** | ● | ox | embryo development, transport, cell motility, others |
|  |  |  | Population | 50.32 | 3, 40 | **>0.001\*** |  |  |  |
|  |  |  | Temp x Pop | 2.95 | 3, 40 | **0.044\*** |  |  |  |
| 6 | 120 | 65.3% | Temperature | 33.60 | 1, 40 | **>0.001\*** | ● | heat |  |
|  |  |  | Population | 8.06 | 3, 40 | **>0.001\*** |  |  |  |
|  |  |  | Temp x Pop | 1.06 | 3, 40 | 0.378 |  |  |  |
| 7 | 119 | 49.5% | Temperature | 17.18 | 1, 40 | **>0.001\*** | ● | ox | translation |
|  |  |  | Population | 44.89 | 3, 40 | **>0.001\*** |  |  |  |
|  |  |  | Temp x Pop | 10.90 | 3, 40 | **>0.001\*** |  |  |  |
| 8 | 98 | 55.8% | Temperature | 9.07 | 1, 40 | **0.004\*** | ● | ox |  |
|  |  |  | Population | 26.99 | 3, 40 | **>0.001\*** |  |  |  |
|  |  |  | Temp x Pop | 3.34 | 3, 40 | **0.029\*** |  |  |  |
| 9 | 77 | 67.9% | Temperature | 13.42 | 1, 40 | **0.001\*** | ● | heat | metabolic<br>process,<br>aging |
|  |  |  | Population | 6.56 | 3, 40 | **0.001\*** |  |  |  |
|  |  |  | Temp x Pop | 0.79 | 3, 40 | 0.506 |  |  |  |
| 10 | 72 | 58.9% | Temperature | 0.01 | 1, 40 | 0.931 | ● | ox, heat |  |
|  |  |  | Population | 51.13 | 3, 40 | **>0.001\*** |  |  |  |
|  |  |  | Temp x Pop | 1.49 | 3, 40 | 0.232 |  |  |  |
| 11 | 60 | 58.8% | Temperature | 27.77 | 1, 40 | **>0.001\*** |  |  | metabolic<br>process |
|  |  |  | Population | 6.25 | 3, 40 | **0.001\*** |  |  |  |
|  |  |  | Temp x Pop | 0.32 | 3, 40 | 0.809 |  |  |  |
| 12 | 49 | 68.6% | Temperature | 24.95 | 1, 40 | **>0.001\*** |  |  |  |
|  |  |  | Population | 8.38 | 3, 40 | **>0.001\*** |  |  |  |
|  |  |  | Temp x Pop | 1.56 | 3, 40 | 0.213 |  |  |  |
| 13 | 32 | 65.5% | Temperature | 1.07 | 1, 40 | 0.306 | ● | ox, heat |  |
|  |  |  | Population | 19.80 | 3, 40 | **>0.001\*** |  |  |  |
|  |  |  | Temp x Pop | 3.84 | 3, 40 | **0.017\*** |  |  |  |
| Unassigned | 9212 | 9.3% | Temperature | 13.96 | 1, 40 | **0.001\*** | ● | ox, heat |  |
|  |  |  | Population | 21.11 | 3, 40 | **>0.001\*** |  |  |  |
|  |  |  | Temp x Pop | 0.59 | 3, 40 | 0.625 |  |  |  |

<sup>a</sup>Significant difference (Tukey's HSD; $p < 0.05$ ) between ancestor and any evolved line

**Table 2.**
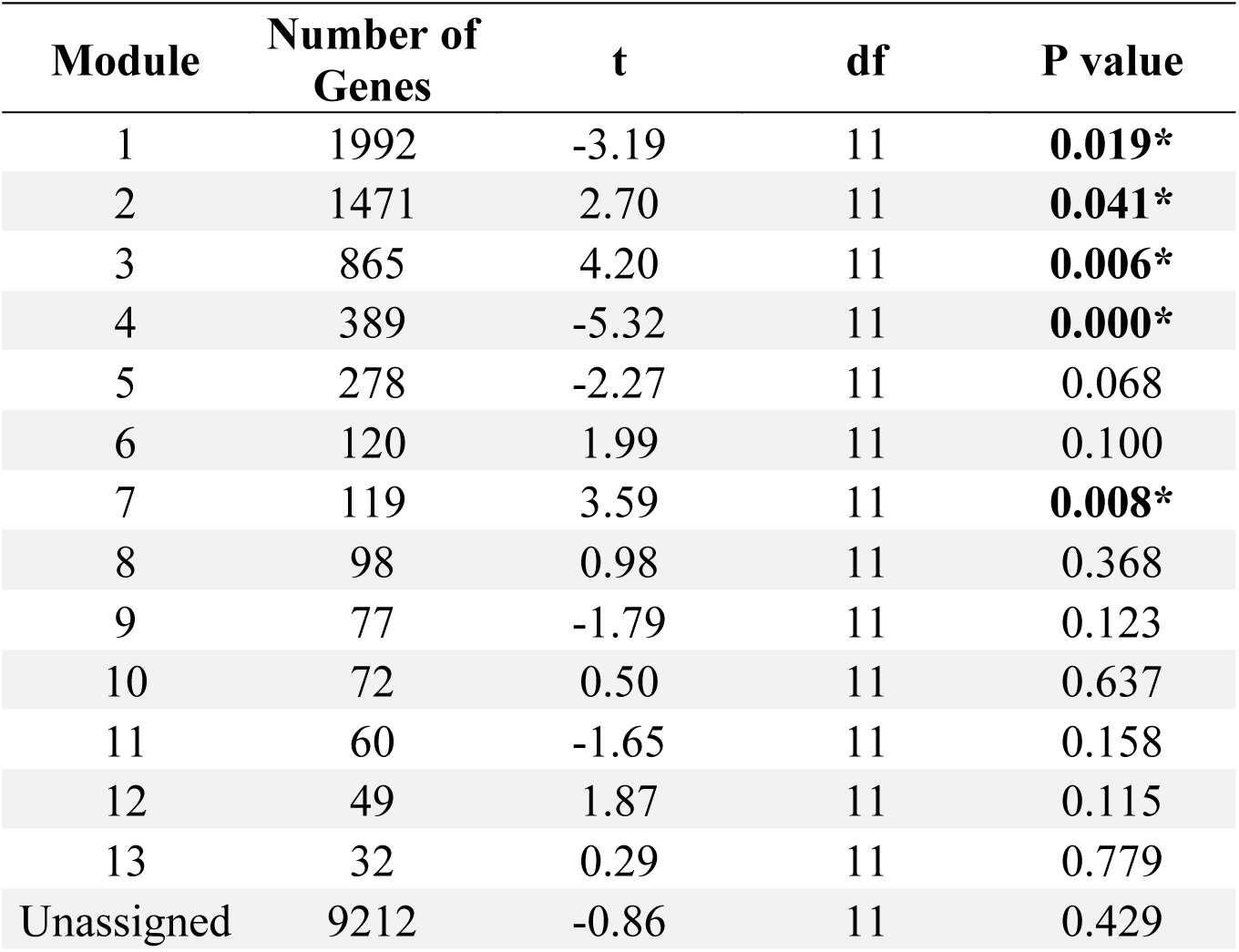
Effect of temperature on module expression in the ancestral population.

## SUPPLEMENTAL FIGURE LEGENDS

**Supplemental Figure S1.** Schematic representation of the transcriptional profiling experiment. Four populations were considered: the ancestor and three experimentally evolved populations selected under different environmental stresses. The induction of transcriptional plasticity was assessed in each line across two different thermal environments.

**Supplemental Figure S2.** Comparison of eigengene expression with overall expression in corresponding module genes. For each module, the first panel shows mean-centered expression values for each gene within the module (light-colored lines), as well as the sample mean and standard deviation for all genes in the module. The second column shows the mean eigengene expression (+/− 1 standard deviation) for each treatment. Percent variance explained by the eigengene (PC1) and principal components 2-5 is shown in the third panel.

**Supplemental Figure S3.** Enrichment of molecular function gene ontology terms in coexpression modules. Red outlines and shading signify that there is significant enrichment of genes mapping to the GO term within the module (FDR<0.05). The intensity of the shading corresponds to the odds ratio for the GO term.

**Supplemental Figure S4.** Enrichment of cellular component gene ontology terms in coexpression modules. Red outlines and shading signify that there is significant enrichment of genes mapping to the GO term within the module (FDR<0.05). The intensity of the shading corresponds to the odds ratio for the GO term.

